# Circulating Platelets Modulate Oligodendrocyte Progenitor Cell Differentiation During Remyelination

**DOI:** 10.1101/2023.09.20.558582

**Authors:** Amber R. Philp, Carolina R. Reyes, Josselyne Mansilla, Amar Sharma, Chao Zhao, Carlos Valenzuela-Krugmann, Khalil S. Rawji, Ginez A. Gonzalez Martinez, Penelope Dimas, Bryan Hinrichsen, César Ulloa-Leal, Amie K. Waller, Diana M. Bessa de Sousa, Maite A. Castro, Ludwig Aigner, Pamela Ehrenfeld, Maria Elena Silva, Ilias Kazanis, Cedric Ghevaert, Robin J.M. Franklin, Francisco J. Rivera

## Abstract

Revealing unknown cues that regulate oligodendrocyte progenitor cell (OPC) function in remyelination is important to optimise the development of regenerative therapies for multiple sclerosis (MS). Platelets are present in chronic non-remyelinated lesions of MS and an increase in circulating platelets has been described in experimental autoimmune encephalomyelitis (EAE) mice, an animal model for MS. However, the contribution of platelets to remyelination remains unexplored. Here we show platelet aggregation in proximity to OPCs in areas of experimental demyelination. Partial depletion of circulating platelets impaired OPC differentiation and remyelination, without altering blood-brain barrier stability and neuroinflammation. Transient exposure to platelets enhanced OPC differentiation *in vitro*, whereas sustained exposure suppressed this effect. In a mouse model of thrombocytosis (*CALR^HET^*), there was a sustained increase in platelet aggregation together with a reduction of newly-generated oligodendrocytes following toxin-induced demyelination. These findings reveal a complex bimodal contribution of platelet to remyelination and provide insights into remyelination failure in MS.

## Introduction

In the central nervous system (CNS), remyelination by newly generated oligodendrocytes is largely mediated by the differentiation of oligodendrocyte progenitor cells (OPCs). In response to demyelination, OPCs proliferate, migrate, and differentiate into remyelinating oligodendrocytes (Franklin & ffrench-Constant, 2008). Although remyelination represents a robust regenerative response to demyelination, it fails during the progress of multiple sclerosis (MS), a CNS autoimmune demyelinating disease (Noseworthy *et al*., 2000). Unravelling the mechanisms that govern remyelination is essential to our understanding of why this important regenerative process fails in MS, as well as in guiding the development of regenerative therapies.

Platelets are small, anucleate cells essential for haemostatic plug formation (Semple *et al*., 2011). Platelets also display tissue-regenerative properties (Nurden, 2011). Several growth factors known to modulate OPCs’ responses to demyelination, such as PDGF and FGF2 (Woodruff *et al*., 2004; Murtie *et al*., 2005; Zhou *et al*., 2006; Clemente *et al*., 2011; Hiratsuka *et al*., 2019), are stored in platelets (Chen *et al*., 2012; Lohmann *et al*., 2012; Schallmoser & Strunk, 2013; Warnke *et al*., 2013). We have previously shown that platelet lysate increases neural stem / progenitor cells (NSPCs) survival, an alternative but infrequent cellular source for mature oligodendrocytes (Kazanis *et al*., 2015). Although this evidence argues in favour of a beneficial contribution of platelets to remyelination, other studies suggest a detrimental role. CD41-expressing platelets and platelet-contained molecules are found in non-remyelinated MS lesions (Lock *et al*., 2002; Han *et al*., 2008; Langer *et al*., 2012; Simon, 2012; Steinman, 2012). Moreover, MS patients show increased levels of circulating platelet microparticles (PMPs) (Marcos-Ramiro *et al*., 2014) and the number of PMPs are indicative of the clinical status of the disease (Saenz-Cuesta *et al*., 2014). Additionally, MS patients display high plasma levels of platelet-specific factors such as, P-selectin and PF4 that correlate with disease course and severity, respectively (Cananzi *et al*., 1987; Kuenz *et al*., 2005). In the animal model for MS, experimental autoimmune encephalomyelitis (EAE), platelet numbers within CNS increase (Sonia D’Souza *et al*., 2018). When platelets were immunodepleted before clinical onset, EAE severity is decreased (Langer *et al*., 2012; Kocovski *et al*., 2019). Here we ask whether circulating platelets regulate OPC function and how this impacts remyelination.

## Results

### Circulating platelets transiently accumulate in response to demyelination and accumulate in close proximity to OPCs

We first assessed the distribution of platelets during remyelination. We created lysolecithin (LPC)-induced demyelinating lesions in the spinal cord white matter of wild type (WT) mice and collected tissue sections at 1-, 3-, 5-, 7-, 10 and 14-days post-lesion (dpl). We observed CD41+ platelet aggregates within and around the lesion early after demyelination (3 dpl) (p-value < 0.01) (Figure 1A and B). However, this was transient as platelet aggregates subsequently decreased until no aggregates were detected at 14 dpl (Figure 1A and B). To assess whether platelet recruitment was specific to demyelination we injected PBS containing DAPI directly into the spinal cord. No signs of demyelination were observed under these conditions and platelet aggregation was minimal at 1- and 3-days post-PBS injection (Figure 1C). We next evaluated the localization of platelets within the lesion. Large platelet aggregates were found within the blood vessels and within the tissue parenchyma at 5 dpl (Figure 1D). Platelets often localised with Olig2^+^ cells around blood vessels, a scaffold used by OPCs for migration (Tsai *et al*., 2016) (Figure 1D).

**Figure 1:**
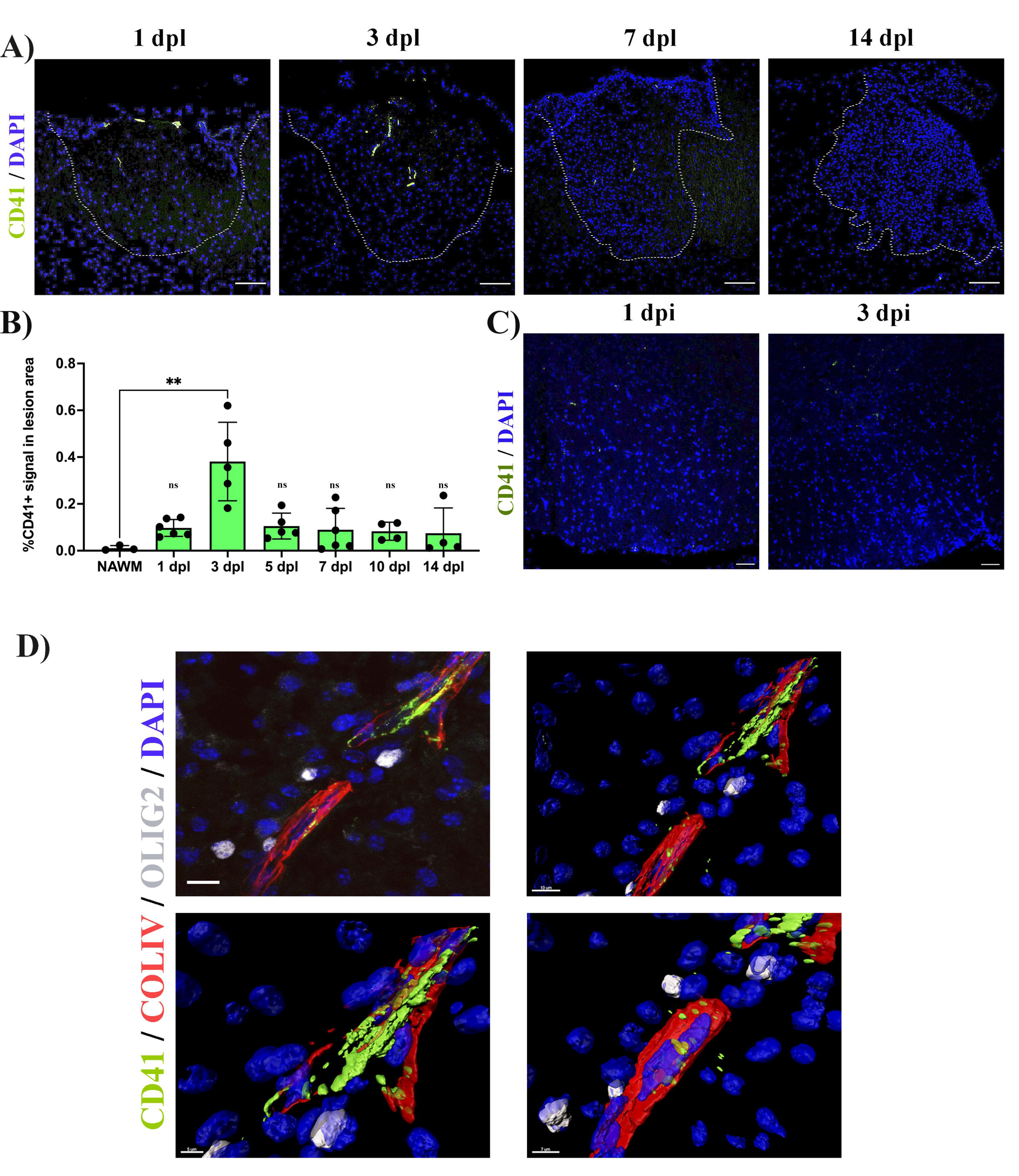
Platelets accumulate in response to demyelination. A) LPC induced demyelinating lesions in spinal cord white matter of WT mice at 1, 3, 7, and 14 dpl, stained for platelets (CD41^+^). Scale bar 100 μm. B) Quantification of CD41^+^ signal within the demyelinated lesion at 1 (*n=6*), 3 (*n=5*), 5 (*n=5*), 7 (*n=6*), 10 (n=4), and 14 dpl (*n=4*), and in NAWM (*n=3*). Scale bar 50 μm. C) Platelet staining (CD41^+^) in spinal cord white matter injected with PBS/DAPI. D) Upper left panel: localization of platelets within blood vessels (ColIV^+^) and in close proximity with OPCs (Olig2^+^) at 5 dpl. Upper right panel: IMARIS 3D projection shows the spatial distribution of platelets. Lower panels: magnification of the IMARIS projection showing platelet aggregation within the blood (left panel) and penetration into the parenchyma (right panel). Data were analysed using one-way ANOVA. Data represent the mean ± SD. *p<0.05.

### Depletion of circulating platelets alters OPC differentiation and remyelination in vivo

To investigate whether circulating platelets modulate OPC function *in vivo*, we used a platelet depletion model (Figure 2A). LPC-induced focal demyelinating lesions were performed in WT mice followed by the administration of anti-CD42b at 3 dpl and every second day to prevent further platelet recruitment (Morodomi *et al*., 2020; de Sousa *et al*., 2023). We first confirmed that this depletion strategy leads to decreased numbers of recruited platelets, with no accumulation in the lesion (p-value < 0.05) (Figure 2B). At 7 dpl, there was no difference in the number of Olig2^+^ cells within the lesion between the platelet depleted and untreated group (Figure 2C, upper panels, and D), indicating that platelets do not alter OPC recruitment in response to demyelination. Through the detection of CC1 expression, a marker that identifies mature oligodendrocytes (Figure 2C, lower panels), we found that platelet depletion significantly decreased the number and percentage of Olig2^+^/CC1^+^ cells compared to untreated mice (p-value ≤ 0.05) (Figure 2E and 2F), indicating that platelet depletion impairs OPC differentiation. Consistently, at 14 dpl we observed a significant decrease in the extent of remyelination (Figure 2G and 2H) and the percentage of remyelinated axons compared to untreated animals (p-value < 0.05) (Figure 2I). Previous studies have shown that decreasing the number of circulating platelets increases blood vessel leakiness (Cloutier *et al*., 2012; Gupta *et al*., 2020). To assess whether impaired OPC differentiation might be due to fibrinogen extravasation (Petersen *et al*., 2017) or enhanced demyelination due to neutrophil infiltration (Ruther *et al*., 2017), we evaluated their presence within the lesion parenchyma after platelet depletion. There were no significant differences between neutrophil (Supplementary Figure 1A and B) and fibrinogen extravasation (Supplementary Figure 1C and D) after platelet depletion at 7 dpl, indicating that remyelination impairment likely derives from low numbers of circulating platelets rather than increased vascular leakiness.

**Figure 2:**
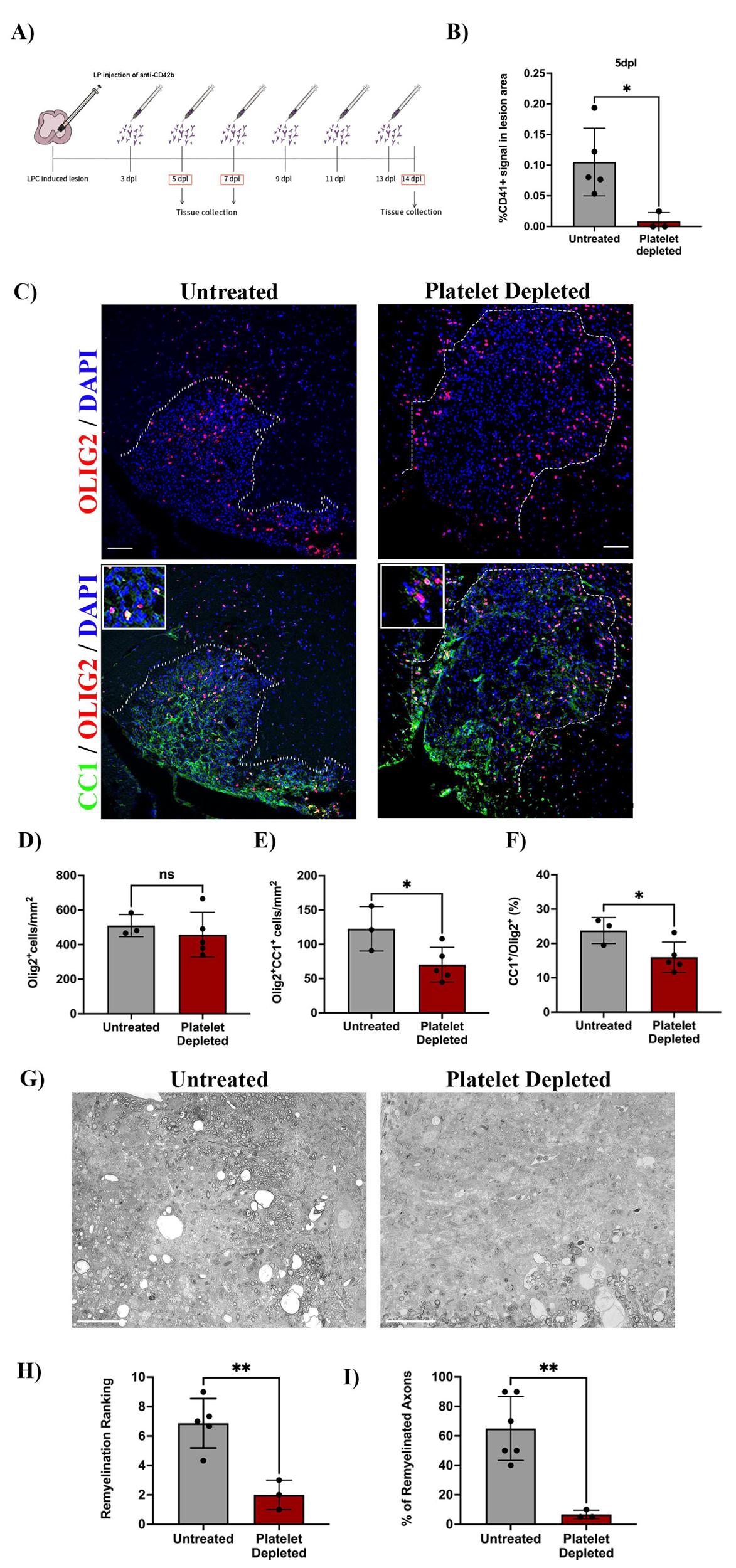
Platelet depletion impairs remyelination *in vivo*. A) Schematic representation of the LPC-induced demyelination model coupled with platelet depletion using anti-CD42b. B) Quantitative analysis of CD41^+^ signal at 5 dpl in untreated *(n=5)* and platelet depleted mice *(n=3)*. C) Representative images of immunofluorescence staining of oligodendroglial lineage cells in untreated and platelet depleted mice at 7 dpl using Olig2^+^ (upper panels) and mature oligodendrocytes using Olig2^+^/CC1^+^ (lower panels). Boxed areas represent high magnification images. (D-F) Quantitative analysis of oligodendroglia at 7 dpl in untreated (*n=3*) and platelet depleted mice (*n=5*). G) Representative images of toluidine blue staining of remyelination in untreated *(n=5)* and platelet depleted mice *(n=3)* at 14 dpl and (H-I) its quantification by relative ranking analysis. Data were analyzed using an Unpaired Student’s t-test or Mann-Whitney U test. Data represent mean ± SD. *p ≤0.05; ns (not significant), p >0.05. Scale bars, 100 μm.

### Depletion of circulating platelets does not alter macrophage/microglia numbers and polarization during remyelination

Blood-borne macrophages and CNS-resident microglia are essential for OPC differentiation during remyelination (Kotter *et al*., 2006; Miron *et al*., 2013). As platelets regulate macrophage function in neuroinflammation (Langer & Chavakis, 2013; Carestia *et al*., 2019; Rolfes *et al*., 2020) and since platelets are located near macrophages/microglia upon demyelination (Supplementary Figure 2A), we evaluated whether platelet depletion affects these cell populations (Supplementary Figure 2B). At 10 dpl, platelet depletion did not alter the total number of IBA-1^+^ (Supplementary Figure 2C), pro-inflammatory IBA-1^+^/CD16/32^+^ (Supplementary Figure 2D) or anti-inflammatory IBA-1^+^/Arg-1^+^ (Supplementary Figure 2E) macrophages/microglia present within the remyelinating lesion. Furthermore, platelet depletion did not influence macrophage/microglia phagocytic activity as no difference in myelin debris clearance, detected by Oil-red O, was observed (Supplementary Figure 2F and G). Therefore, circulating platelets likely impact OPC differentiation without interfering with macrophage/microglia numbers/polarization during remyelination.

### Transient in vitro exposure to platelets enhances OPC differentiation

To confirm whether transient platelet exposure directly enhances OPC differentiation, OPCs were briefly exposed to washed platelets (WP) for 3 days (pulse) and differentiation was assessed 3 days after WP withdrawal. OPCs briefly exposed to 10% WP exhibited a significant increase in the percentage of Olig2^+^/MBP^+^ mature oligodendrocytes compared to the vehicle treated control (p-value < 0.0001) (Figure 3A and B), indicating that transient contact to platelets directly promotes OPC differentiation. Similar increases in the proportion of Olig2^+^/MBP^+^ mature oligodendrocytes were observed when OPCs were transiently exposed to 1% platelet lysate (PL) compared to vehicle-treated control, indicating that this effect is, at least in part, mediated through platelet-contained factors and direct cell-cell contact is not essential (p-value < 0.05) (Figure 3C - D).

**Figure 3:**
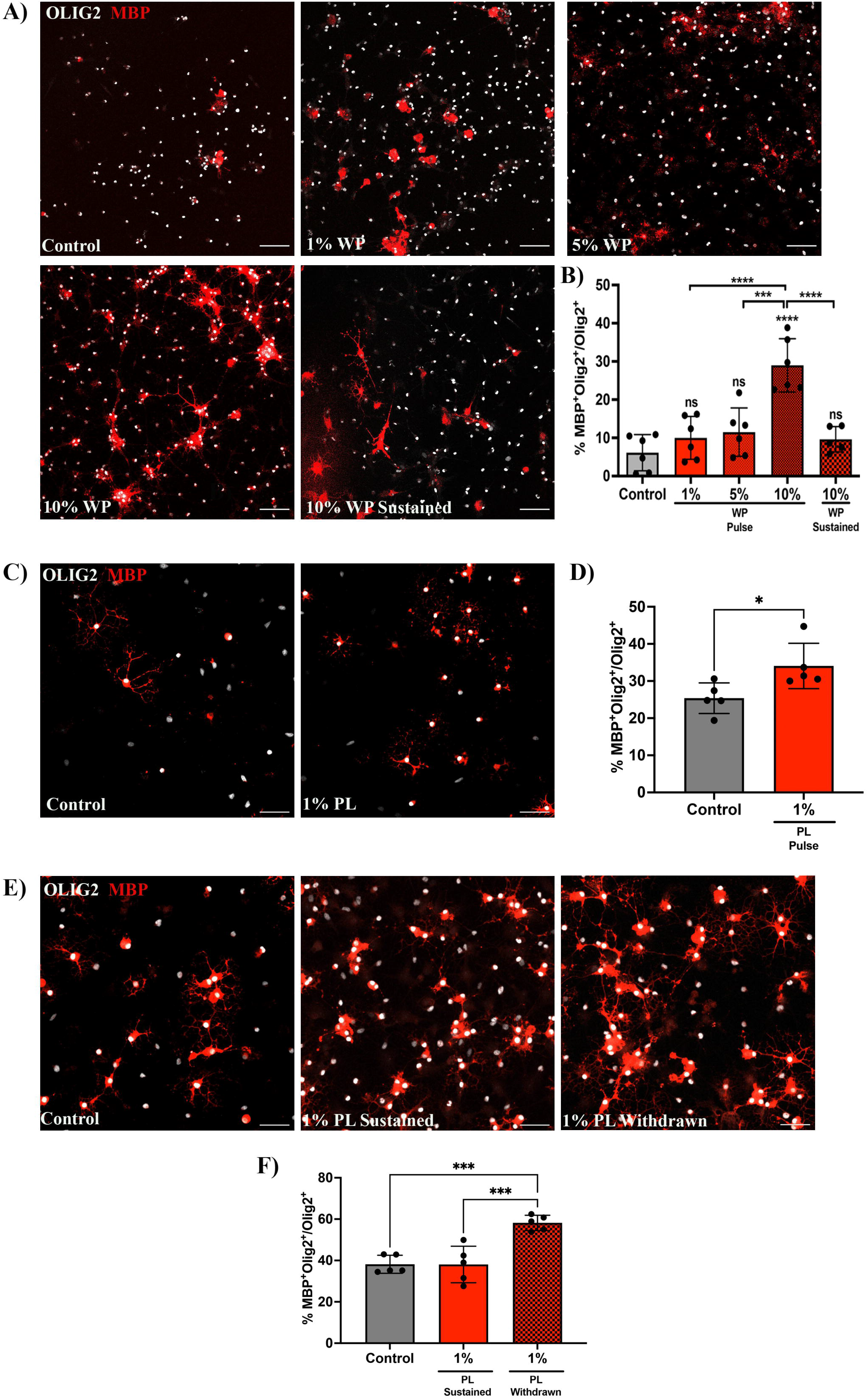
Prolonged exposure to platelets suppresses their ability to enhance OPC differentiation. A) Representative fluorescence images of OPCs co-cultured with 1 (*n=6*), 5 (*n=6*), and 10% (*n=6*) washed platelets (WP) for 3 days *in vitro* (DIV), followed by WP removal for an additional 3 DIV (Pulse). Additionally, OPCs were co-cultured in the presence of 10% WP for 6DIV (*n=5*) (Sustained). Vehicle treated OPCs represents the control condition (*n=6*). B) Graph represents the percentage of Olig2^+^MBP^+^ oligodendrocytes within the total Olig2 population (quantitative analysis of OPC differentiation). C) Representative images of OPCs exposed to 1% platelet lysate (PL) (*n=5*) for 6 DIV. Vehicle treated OPCs represents the control condition (*n=5*). D) Graph represents the quantitative analysis of OPC differentiation as in B. E) Representative images of OPCs exposed to either PL for 9 DIV (Sustained) (*n=5*) or 6DIV with PL followed by its removal for an additional 3 more DIV (Withdrawn) (*n=5*). Vehicle treated OPCs represents the control condition (*n=5*). F) Graph shows the quantitative analysis of OPC differentiation as in B and D. Data were analyzed using One-way Anova followed by Tukey’s post-hoc test or an un-paired t-test. Data represent the mean ± SD. *** p ≤0.001; **** p ≤0.0001; ns (not significant), p >0.05. Scale bars, 50 mm.

### Sustained increase in circulating platelets hampers OPC differentiation during remyelination

Chronically-demyelinated MS lesions have been reported to contain a substantial number of platelets and their derived molecules (Lock *et al*., 2002; Han *et al*., 2008; Langer *et al*., 2012; Simon, 2012; Steinman, 2012). To explore the effects of prolonged platelet exposure on OPC differentiation we conducted experiments with sustained exposure to 10% WP. Contrary to the 3-day pulse-based exposure, 6-days of sustained exposure to 10% WP suppressed the ability of platelets to enhance OPC differentiation (Figure 3A and B). Similar findings were observed upon 9-days of sustained exposure to 1% PL (Figure 3E and F), indicating effects mediated by platelet-contained factors. To test whether this effect is reversible, PL was withdrawn upon 6 days of sustained exposure, and OPC differentiation was evaluated 3 days later. Interestingly, PL withdrawal rescued the capability of platelets to enhance OPC differentiation when compared to the vehicle treated control and the sustained condition (p-value < 0.0006) (Figure 3E - F).

To assess whether a permanent increase of circulating platelets may hamper OPC differentiation during remyelination, we used a conditional mouse knock-in model carrying a mutation within the calreticulin gene in a heterozygous fashion controlled by the VaV hematopoietic promoter, resulting in sustained thrombocytosis (2 to 3 times more circulating platelets) without alterations in other cell lineages (Li *et al*., 2018). We induced a demyelinating lesion by LPC injection in the spinal cord white matter of CALR^HET^ mice and evaluated platelet recruitment and OPC differentiation. As expected, CALR^HET^ mice showed increased levels of circulating platelets (p-value < 0.01) (Figure 4B) as well as a higher number of recruited platelets into the lesion (p-value < 0.05) (Figure 4A and C). At 10 dpl, CALR mice displayed a reduced number of mature Olig2^+^/CC1^+^ oligodendrocytes (Figure 4D and F) and a significant decrease in the percentage of differentiated OPCs (p-value < 0.05) (Figure 4G) compared to WT mice, without alterations in the total number of Olig2^+^ cells (Figure 4E). Additionally, we observed a negative correlation between the number of circulating platelets in CALR^HET^ mice with the number of mature oligodendrocytes (r= -0.75, p-value < 0.01) (Figure 4H). Similar to the platelet depleted model, effects on OPC differentiation are not mediated by inflammation, as CALR^HET^ mice showed no alterations in macrophage/microglia numbers/polarization during remyelination (Supplementary Figure 2B-E). These findings indicate that sustained exposure to platelets directly hampers OPC differentiation during remyelination.

**Figure 4:**
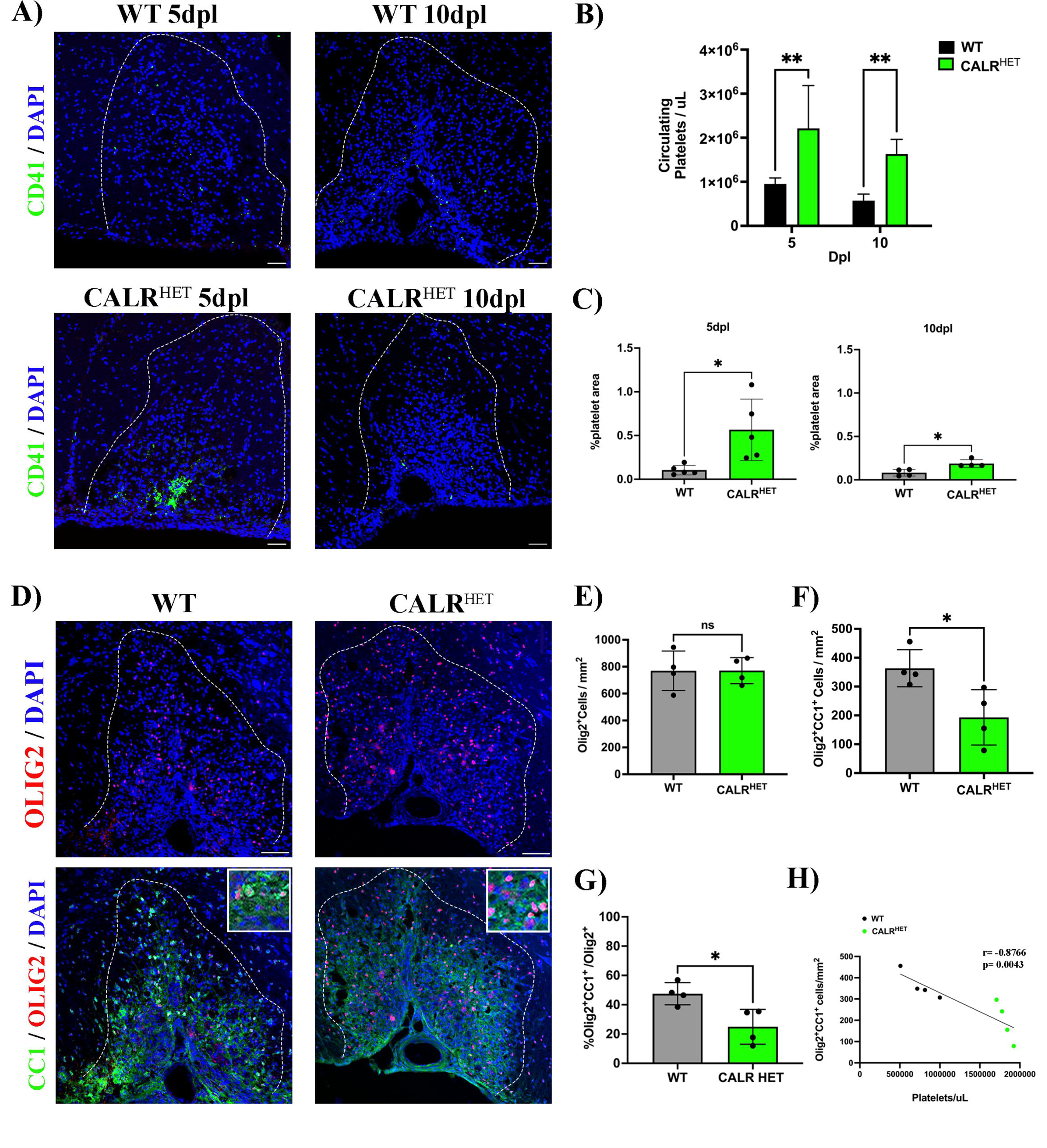
A sustained increase in circulating platelets impairs remyelination *in-vivo*. A) Representative fluorescence images of platelets (CD41^+^) in LPC induced demyelinating lesions of spinal cord white matter of WT and CALR^HET^ mice at 5 and 10 dpl B) Quantification of circulating platelets in WT vs CALR^HET^ mice at 5 (*n=4 & n=5, respectively*), 10 ((*n=5 & n=6, respectively*), and 14 dpl ((*n=4 & n=6, respectively*). C) Quantification of CD41^+^ signal in demyelinated lesions of WT vs CALR^HET^ mice at 5 dpl (*n=5 & n=5*, respectively) and 10 dpl (*n=4 & n=4*, respectively). D) Representative immunofluorescence staining of oligodendroglial lineage cells in untreated and platelet depleted mice at 10 dpl using Olig2^+^ (upper panels) and mature oligodendrocytes using Olig2^+^/CC1^+^ (lower panels) (*n=4*). Scale bar 100 μm. (E-G) Quantitative analysis of oligodendroglia at 10 dpl. H) Correlation between the circulating platelet number with the number of Olig2^+^/CC1^+^ cells within the demyelinated lesion. Data were analyzed using an unpaired t-test or Spearman’s rank correlation analysis. Data represent the mean ± SD. *p ≤0.05; ** p ≤0.001; ns (not significant), p >0.05.

## Discussion

In conclusion, our study reveals that in response to myelin damage platelets transiently accumulate within the vascular niche and locate near OPCs. While transient contact to platelets support OPC differentiation, long lasting exposure to elevated numbers of circulating platelets hampers the generation of oligodendrocytes during remyelination. These findings argue in favour of a beneficial physiological role of platelets in remyelination. However, we also highlight that sustained increased platelet counts, as occurs in MS-related conditions, negatively alter OPC function and contribute to remyelination failure in MS.

Although there is a need to reveal the underlying mechanism(s) by which platelets exert a bimodal action on OPC differentiation, our findings indicate that platelet-contained factors contribute to this effect. This study shows that the regeneration of oligodendrocytes rests on the transient vs sustained presence of platelets within demyelinated lesions. Platelet accumulation in MS lesions may result from blood-brain barrier damage (Broman, 1964; Zlokovic, 2008) and/or a clearance failure, but changes in their adhesiveness (Sanders *et al*., 1968) and hyperactivity observed during MS (Sheremata *et al*., 2008) may contribute to such scenario. Strategies that restore platelet function, spatially and temporally, represent a future step for developing regenerative therapies in MS.

## Materials and methods

### Animals

All animal work at University of Cambridge complied with the requirements and regulations of the United Kingdom Home Office (Project Licenses PCOCOF291 and P667BD734). All the experiments at Universidad Austral de Chile were conducted in agreement with the Chilean Government’s Manual of Bioethics and Biosafety (CONICYT: The Chilean Commission of Scientific and Technological Research, Santiago, Chile) and according to the guidelines established by the Animal Protection Committee of the Universidad Austral de Chile (UACh). The animal study was reviewed and approved by the *Comité Institucional de Cuidado y Uso de Animales (CICUA)-UACh* (Report Number # 394/2020). All the experiments at University of Helsinki followed the guidelines posed by the Academy of Finland and the University of Helsinki on research ethics and integrity (under Internal License KEK23-022) and accordingly to the National Animal Ethics Committee of Finland (ELLA). Mice and rats had access to food and water *ad libitum* and were exposed to a 12-hour light cycle.

### Human subjects

Human platelets were obtained from blood samples of healthy volunteers who signed a consent form before sampling. All procedures were approved by the *Comité Ético y Científico del Servicio de Salud de Valdivia* (CEC-SVS) (ORD N° 510) to carry experiments at Universidad Austral de Chile and by the Ethical Committee of the University of Cambridge to perform experiments at this institution. The blood donors at Cambridge were approved by the human biology research ethics committee (reference number: HBREC.2018.13.).

### Focal demyelination lesions

A focal demyelinating lesion was induced in C57BL/6 and CALR^HET^ mice between 2-4 months of age. Animals were anesthetized using Isoflurane/Oxygen (2-2.5%/1000 ml/min O_2_) and buprenorphine (0.05 mg/kg) was injected subcutaneously immediately before surgery. Local Lysolecithin-driven demyelination in mice was induced as previously described in (Fancy *et al*., 2009). Briefly, the spinal cord was exposed between two vertebrae of the thoracic column and demyelination was induced by injecting 1uL of 1% lysolecithin (L-lysophosphatidylcholine, Sigma) into the ventral funiculus at a rate of approximately 0.5 µl/min^−1^. The incision was then sutured, and the animal was left to recover in a thermally controlled chamber. Animals were monitored for 72 hours after surgery. Any signs of pain, dragging of limbs, or weight loss of more than 15% of pre-surgery weight, resulted in cessation of the experiment. Mice were sacrificed at 1, 3, 5, 7, 10, and 14 dpl by transcardial perfusion of 4% PFA or glutaraldehyde under terminal anaesthesia.

### Platelet Depletion

For platelet depletion, mice received an intraperitoneal injection (IP) of 0.6µg/g of antiCD42b (Emfret Analytics) (Evans *et al*., 2021), diluted in saline solution, at 3 dpl, followed by IP injections every 48 hours until the end of the experiment period. The effectiveness of platelet depletion was confirmed by measuring the number of circulating platelets using a VetAnalyzer (scil Vet abc Plus). Mice with a circulating platelet number below 200,000 platelets/uL were considered successfully depleted.

### Preparation of washed platelets and platelet lysate

Washed platelets (WP) were prepared as described (Cazenave *et al*., 2004). Briefly, human blood samples were taken from the median cubital vein and collected in sodium citrate followed by centrifugation for 20 mins at 120x g to separate the red blood cells from the plasma. Plasma was collected and centrifuged at 1400x g to pellet platelets. Plasma was removed without disrupting the platelet pellet. PGI_2_ and sodium citrate were carefully added, followed by resuspension in Tyrode’s buffer. Platelet number was quantified using a Vet Analyzer and adjusted to a concentration of 1,000,000 platelets/uL. WP were used fresh, meanwhile for the platelet lysate (PL) preparation, the suspension underwent two freeze-thaw overnight cycles. Platelet fragments were then eliminated by centrifugation at 4000x g for 15 minutes and the supernatant was collected and stored at - 20°C.

### Primary OPC cultures

Rat OPCs were isolated as described by (Neumann *et al*., 2019). Cells were then seeded onto glass plates pre-coated with Poly-D-Lysine (PDL) in 24-well plates, with a seeding density of 7,000 cells for differentiation assays and 10,000 cells for proliferation assays. For differentiation conditions, T3 was added to the culture media. All experimental conditions were replicated using two independent technical replicates. OPCs were either subjected to various concentrations of washed platelets (1%, 5%, and 10%) or to 1% of platelet lysate of the final volume.

### Histology and Immunofluorescence

After transcardial perfusion with 4% PFA, tissue was post-fixed overnight in 4% PFA at 4°C. After fixation, spinal cords were left in 30% sucrose overnight. Tissue was then embedded in OCT and cut in 15um transverse sections on a Leica Cryostat. Samples were stored at - 80°C until use.

For immunofluorescence staining of tissues, samples were left to thaw for 30 mins and washed with PBS. Samples were blocked for 1 hour, using a blocking solution that contained; 10% horse serum, 1% bovine serum albumin, 0.1% cold fish gelatin, 0.1% Triton X-100, and 0.05% Tween 20, diluted in PBS. After blocking, samples were incubated overnight at 4°C with primary antibody diluted in PBS containing 1% bovine serum albumin, 0.1% cold fish gelatin, and 0.5% Triton X-100. The following primary antibodies were used: rat anti-CD41 (1:200 Abcam), rat anti-CD16/32 (1:200, Santa Cruz), rabbit anti-IBA1 (1:500, WAKO), rat anti-CD31 (1.300 BD Biosciences), rabbit anti-Olig2 (1:200, Abcam), rabbit anti-Ki67 (1:100, Abcam), goat anti-Arg1 (1:200, Santa Cruz), mouse anti-CC1 (1:1000, Calbiochem), rat anti-NIMP-R14 (1:200, Abcam). Samples were washed 3 times for 5mins in PBS. After washing, samples were incubated with secondary antibody and DAPI for 1 hour, diluted in the same solution as the primary antibody. Samples were washed 3 times for 5 mins in PBS. Samples were mounted with Fluromount. All secondary antibodies were diluted 1:500. For imaging of spinal cord tissue, the entire lesion area was imaged for 5 technical replicates.

For immunofluorescence staining of cell cultures, samples were initially washed three times with PBS for 5 mins after fixation. The cells were then blocked with 10% Donkey Serum (DKS) in PBS for 1 hour, followed by incubation with the primary antibody overnight, diluted in the same blocking solution. The following primary antibodies were utilized: Rat Anti-Myelin Basic Protein (MBP; 1:500, Biorad) and Rabbit Anti-Oligodendrocyte transcription factor 2 (Olig2; 1:500, Abcam). The cells were then washed three times with PBS 1x for 5 mins, followed by incubation with secondary antibodies, diluted in the blocking solution, for 1 hour. The cells were washed three more times with PBS 1x for 5 mins.

Images were captured using a Leica SP8 Laser Confocal, a Zeiss LSM 980 Confocal or an Olympus IX81FV1000. For cell culture imaging, 8-10 photos per well were quantified for each well using an automated macro in ImageJ/Fiji. For in-vivo imaging, 3-5 photos were quantified per animal by a blinded observer. For tissue image analysis and 3D reconstruction of platelet localisation, ImageJ/Fiji (version 2.1.0/1.53 h) and Imaris (Bitplane, version 9.3.1, and 9.9.0) were used.

### Oil-red O Staining

To analyse myelin debris clearance, tissue sections were stained with Oil-red O as previously described by (Kotter *et al*., 2005). Briefly, sections were stained with freshly prepared Oil-red O and incubated at 37 degrees for 30 mins. Slides were washed and mounted using an aqueous mounting medium. Image J was used to threshold and quantify Oil-red O images.

### Remyelination Ranking Analysis

For remyelination studies, tissue was fixed with 4% glutaraldehyde and embedded in resin. Semi-thin sections of the lesion were cut and stained with Toluidine Blue. Three blinded observers ranked the level of remyelination for each biological individual, giving the most remyelinated individual the highest score, and the individual with the lowest degree of remyelination the lowest. The average for each animal was calculated from the three independent observer rankings.

### Statistical Analysis

Statistical analysis was performed using GraphPad Prism 8. The distribution of data were first tested using a Shapiro-Wilks test. One-way ANOVA or a Kruskal Wallis one-way analysis, with the corresponding post-hoc test, were used to compared multiple groups, and a Mann-Whiteney U-test or an unpaired t-test was used to compare between groups. P-values were represented as *≤0.05, **≤0.01, ***≤0.001, ***≤0.0001.

## Funding

Authors would like to thank the funding support from *Agencia Nacional de Investigación y Desarrollo* (ANID, Chile)-FONDECYT Program Regular Grant Number 1201706 and 1161787 (both to F.J.R.), ANID-PCI Program Grant N° REDES170233 (to F.J.R.) and N° REDES180139 (to M.A.C), ANID-National Doctoral Fellowship N° 21170732 (to A.R.P) and N° 21211727 (to C.R.R). In addition, the authors would like to thank the PROFI 6 N° 336234 of the Research Council of Finland.

## Acknowledgments

Special acknowledgments to the *Comité Ético y Científico del Servicio de Salud de Valdivia* (CEC-SVS) for ethical guidance and approval (ORD N° 510). The authors would like to thank the Laboratory of Chronobiology, UACh (led by Dr. Claudia Torres-Farfán) for supporting animal experimentation. We would also like to thank Dr. Alerie Guzman de la Fuente for the Fiji/ImageJ macro for *in-vitro* quantification.

## Disclosure statement

The authors declare that there are no conflicts of interest.

## Figure Legends

**Supplementary Figure 1: Platelet depletion does not alter BBB permeability.** A) Representative immunofluorescence images of neutrophils (NIMP-R14^+^) and B) number of neutrophils in LPC-induced white matter spinal cord lesion at 7dp in untreated (*n=3*) and platelet depleted mice (*n=5*). C) Representative immunofluorescence images of Fibrinogen and D) quantification of Fibrinogen signal within the demyelinated lesion at 7 dpl in untreated and platelet depleted mice (*n=4*). Scale bar 50 μm. Data were analyzed using an unpaired t-test. Data represent the mean ± SD. ns (not significant), p >0.05.

**Supplementary Figure 2: Changes in circulating platelet numbers does not alter the macrophage/microglia population during remyelination.** A) Platelets (CD41^+^) are located in close proximity to the macrophage/microglia population (IBA1^+^) at 5 dpl. Scale bar 50 um. B) Total macrophage/microglia population (IBA-1^+^), M1 (CD16/32^+^) and M2 (Arg-1^+^) macrophage/microglia populations at 10 dpl and (C-E) their respective quantitative analysis in untreated (*n=6*), platelet depleted (*n=3*), and CALR^HET^ mice (*n=4*). F) Representative oil-red staining of myelin debris at 10 dpl. G) Quantification of Oil-red staining in untreated (*n=6*) and platelet depleted mice (*n=3*). Scale bar 100μm. Data were analyzed using an unpaired t-test. Data represent the mean ± SD. ns (not significant), p >0.05.

## Notes

### Competing Interest Statement

The authors have declared no competing interest.

### Summary of Updates

Few changes in the text (Material and Methods, Results section and Figure legend 3).

## References

Broman, T. (1964) Blood-Brain Barrier Damage in Multiple Sclerosis Supravital Test-Observations. Acta Neurol Scand Suppl, 40, SUPPL 10:21–14.

Cananzi, A.R., Ferro-Milone, F., Grigoletto, F., Toldo, M., Meneghini, F., Bortolon, F. & D’Andrea, G. (1987) Relevance of platelet factor four (PF4) plasma levels in multiple sclerosis. Acta Neurol Scand, 76, 79–85.

Carestia, A., Mena, H.A., Olexen, C.M., Ortiz Wilczynski, J.M., Negrotto, S., Errasti, A.E., Gomez, R.M., Jenne, C.N., Carrera Silva, E.A. & Schattner, M. (2019) Platelets Promote Macrophage Polarization toward Pro-inflammatory Phenotype and Increase Survival of Septic Mice. Cell reports, 28, 896–908 e895.

Cazenave, J.P., Ohlmann, P., Cassel, D., Eckly, A., Hechler, B. & Gachet, C. (2004) Preparation of washed platelet suspensions from human and rodent blood. Methods in molecular biology, 272, 13–28.

Chen, B., Sun, H.H., Wang, H.G., Kong, H., Chen, F.M. & Yu, Q. (2012) The effects of human platelet lysate on dental pulp stem cells derived from impacted human third molars. Biomaterials, 33, 5023–5035.

Clemente, D., Ortega, M.C., Arenzana, F.J. & de Castro, F. (2011) FGF-2 and Anosmin-1 are selectively expressed in different types of multiple sclerosis lesions. The Journal of neuroscience : the official journal of the Society for Neuroscience, 31, 14899–14909.

Cloutier, N., Pare, A., Farndale, R.W., Schumacher, H.R., Nigrovic, P.A., Lacroix, S. & Boilard, E. (2012) Platelets can enhance vascular permeability. Blood, 120, 1334–1343.

de Sousa, D.M.B., Benedetti, A., Altendorfer, B., Mrowetz, H., Unger, M.S., Schallmoser, K., Aigner, L. & Kniewallner, K.M. (2023) Immune-mediated platelet depletion augments Alzheimer’s disease neuropathological hallmarks in APP-PS1 mice. Aging (Albany NY), 15, 630–649.

Evans, A.L., Dalby, A., Foster, H.R., Howard, D., Waller, A.K., Taimoor, M., Lawrence, M., Mookerjee, S., Lehmann, M., Burton, A., Valdez, J., Thon, J., Italiano, J., Moreau, T. & Ghevaert, C. (2021) Transfer to the clinic: refining forward programming of hPSCs to megakaryocytes for platelet production in bioreactors. Blood Adv, 5, 1977–1990.

Fancy, S.P., Baranzini, S.E., Zhao, C., Yuk, D.I., Irvine, K.A., Kaing, S., Sanai, N., Franklin, R.J. & Rowitch, D.H. (2009) Dysregulation of the Wnt pathway inhibits timely myelination and remyelination in the mammalian CNS. Genes Dev, 23, 1571–1585.

Franklin, R.J. & ffrench-Constant, C. (2008) Remyelination in the CNS: from biology to therapy. Nature reviews. Neuroscience, 9, 839–855.

Gupta, S., Konradt, C., Corken, A., Ware, J., Nieswandt, B., Di Paola, J., Yu, M., Wang, D., Nieman, M.T., Whiteheart, S.W. & Brass, L.F. (2020) Hemostasis vs. homeostasis: Platelets are essential for preserving vascular barrier function in the absence of injury or inflammation. Proc Natl Acad Sci U S A, 117, 24316–24325.

Han, M.H., Hwang, S.I., Roy, D.B., Lundgren, D.H., Price, J.V., Ousman, S.S., Fernald, G.H., Gerlitz, B., Robinson, W.H., Baranzini, S.E., Grinnell, B.W., Raine, C.S., Sobel, R.A., Han, D.K. & Steinman, L. (2008) Proteomic analysis of active multiple sclerosis lesions reveals therapeutic targets. Nature, 451, 1076–1081.

Hiratsuka, D., Kurganov, E., Furube, E., Morita, M. & Miyata, S. (2019) VEGF- and PDGF-dependent proliferation of oligodendrocyte progenitor cells in the medulla oblongata after LPC-induced focal demyelination. Journal of neuroimmunology, 332, 176–186.

Kazanis, I., Feichtner, M., Lange, S., Rotheneichner, P., Hainzl, S., Oller, M., Schallmoser, K., Rohde, E., Reitsamer, H.A., Couillard-Despres, S., Bauer, H.C., Franklin, R.J., Aigner, L. & Rivera, F.J. (2015) Lesion-induced accumulation of platelets promotes survival of adult neural stem / progenitor cells. Experimental neurology, 269, 75–89.

Kocovski, P., Jiang, X., D’Souza, C.S., Li, Z., Dang, P.T., Wang, X., Chen, W., Peter, K., Hale, M.W. & Orian, J.M. (2019) Platelet Depletion is Effective in Ameliorating Anxiety-Like Behavior and Reducing the Pro-Inflammatory Environment in the Hippocampus in Murine Experimental Autoimmune Encephalomyelitis. J Clin Med, 8.

Kotter, M.R., Li, W.W., Zhao, C. & Franklin, R.J. (2006) Myelin impairs CNS remyelination by inhibiting oligodendrocyte precursor cell differentiation. The Journal of neuroscience : the official journal of the Society for Neuroscience, 26, 328–332.

Kotter, M.R., Zhao, C., van Rooijen, N. & Franklin, R.J. (2005) Macrophage-depletion induced impairment of experimental CNS remyelination is associated with a reduced oligodendrocyte progenitor cell response and altered growth factor expression. Neurobiology of disease, 18, 166–175.

Kuenz, B., Lutterotti, A., Khalil, M., Ehling, R., Gneiss, C., Deisenhammer, F., Reindl, M. & Berger, T. (2005) Plasma levels of soluble adhesion molecules sPECAM-1, sP-selectin and sE-selectin are associated with relapsing-remitting disease course of multiple sclerosis. Journal of neuroimmunology, 167, 143–149.

Langer, H.F. & Chavakis, T. (2013) Platelets and neurovascular inflammation. Thrombosis and haemostasis, 110.

Langer, H.F., Choi, E.Y., Zhou, H., Schleicher, R., Chung, K.J., Tang, Z., Gobel, K., Bdeir, K., Chatzigeorgiou, A., Wong, C., Bhatia, S., Kruhlak, M.J., Rose, J.W., Burns, J.B., Hill, K.E., Qu, H., Zhang, Y., Lehrmann, E., Becker, K.G., Wang, Y., Simon, D.I., Nieswandt, B., Lambris, J.D., Li, X., Meuth, S.G., Kubes, P. & Chavakis, T. (2012) Platelets contribute to the pathogenesis of experimental autoimmune encephalomyelitis. Circulation research, 110, 1202–1210.

Li, J., Prins, D., Park, H.J., Grinfeld, J., Gonzalez-Arias, C., Loughran, S., Dovey, O.M., Klampfl, T., Bennett, C., Hamilton, T.L., Pask, D.C., Sneade, R., Williams, M., Aungier, J., Ghevaert, C., Vassiliou, G.S., Kent, D.G. & Green, A.R. (2018) Mutant calreticulin knockin mice develop thrombocytosis and myelofibrosis without a stem cell self-renewal advantage. Blood, 131, 649–661.

Lock, C., Hermans, G., Pedotti, R., Brendolan, A., Schadt, E., Garren, H., Langer-Gould, A., Strober, S., Cannella, B., Allard, J., Klonowski, P., Austin, A., Lad, N., Kaminski, N., Galli, S.J., Oksenberg, J.R., Raine, C.S., Heller, R. & Steinman, L. (2002) Gene-microarray analysis of multiple sclerosis lesions yields new targets validated in autoimmune encephalomyelitis. Nature medicine, 8, 500–508.

Lohmann, M., Walenda, G., Hemeda, H., Joussen, S., Drescher, W., Jockenhoevel, S., Hutschenreuter, G., Zenke, M. & Wagner, W. (2012) Donor age of human platelet lysate affects proliferation and differentiation of mesenchymal stem cells. PloS one, 7, e37839.

Marcos-Ramiro, B., Oliva Nacarino, P., Serrano-Pertierra, E., Blanco-Gelaz, M.A., Weksler, B.B., Romero, I.A., Couraud, P.O., Tunon, A., Lopez-Larrea, C., Millan, J. & Cernuda-Morollon, E. (2014) Microparticles in multiple sclerosis and clinically isolated syndrome: effect on endothelial barrier function. BMC Neurosci, 15, 110.

Miron, V.E., Boyd, A., Zhao, J.W., Yuen, T.J., Ruckh, J.M., Shadrach, J.L., van Wijngaarden, P., Wagers, A.J., Williams, A., Franklin, R.J. & ffrench-Constant, C. (2013) M2 microglia and macrophages drive oligodendrocyte differentiation during CNS remyelination. Nature neuroscience, 16, 1211–1218.

Morodomi, Y., Kanaji, S., Won, E., Ruggeri, Z.M. & Kanaji, T. (2020) Mechanisms of anti-GPIbalpha antibody-induced thrombocytopenia in mice. Blood, 135, 2292–2301.

Murtie, J.C., Zhou, Y.X., Le, T.Q., Vana, A.C. & Armstrong, R.C. (2005) PDGF and FGF2 pathways regulate distinct oligodendrocyte lineage responses in experimental demyelination with spontaneous remyelination. Neurobiology of disease, 19, 171–182.

Neumann, B., Baror, R., Zhao, C., Segel, M., Dietmann, S., Rawji, K.S., Foerster, S., McClain, C.R., Chalut, K., van Wijngaarden, P. & Franklin, R.J.M. (2019) Metformin Restores CNS Remyelination Capacity by Rejuvenating Aged Stem Cells. Cell stem cell, 25, 473–485 e478.

Noseworthy, J.H., Lucchinetti, C., Rodriguez, M. & Weinshenker, B.G. (2000) Multiple sclerosis. The New England journal of medicine, 343, 938–952.

Nurden, A.T. (2011) Platelets, inflammation and tissue regeneration. Thrombosis and haemostasis, 105 **Suppl 1**, S13–33.

Petersen, M.A., Ryu, J.K., Chang, K.J., Etxeberria, A., Bardehle, S., Mendiola, A.S., Kamau-Devers, W., Fancy, S.P.J., Thor, A., Bushong, E.A., Baeza-Raja, B., Syme, C.A., Wu, M.D., Rios Coronado, P.E., Meyer-Franke, A., Yahn, S., Pous, L., Lee, J.K., Schachtrup, C., Lassmann, H., Huang, E.J., Han, M.H., Absinta, M., Reich, D.S., Ellisman, M.H., Rowitch, D.H., Chan, J.R. & Akassoglou, K. (2017) Fibrinogen Activates BMP Signaling in Oligodendrocyte Progenitor Cells and Inhibits Remyelination after Vascular Damage. Neuron, 96, 1003–1012 e1007.

Rolfes, V., Ribeiro, L.S., Hawwari, I., Bottcher, L., Rosero, N., Maasewerd, S., Santos, M.L.S., Prochnicki, T., Silva, C.M.S., Wanderley, C.W.S., Rothe, M., Schmidt, S.V., Stunden, H.J., Bertheloot, D., Rivas, M.N., Fontes, C.J., Carvalho, L.H., Cunha, F.Q., Latz, E., Arditi, M. & Franklin, B.S. (2020) Platelets Fuel the Inflammasome Activation of Innate Immune Cells. Cell reports, 31, 107615.

Ruther, B.J., Scheld, M., Dreymueller, D., Clarner, T., Kress, E., Brandenburg, L.O., Swartenbroekx, T., Hoornaert, C., Ponsaerts, P., Fallier-Becker, P., Beyer, C., Rohr, S.O., Schmitz, C., Chrzanowski, U., Hochstrasser, T., Nyamoya, S. & Kipp, M. (2017) Combination of cuprizone and experimental autoimmune encephalomyelitis to study inflammatory brain lesion formation and progression. Glia, 65, 1900–1913.

Saenz-Cuesta, M., Irizar, H., Castillo-Trivino, T., Munoz-Culla, M., Osorio-Querejeta, I., Prada, A., Sepulveda, L., Lopez-Mato, M.P., Lopez de Munain, A., Comabella, M., Villar, L.M., Olascoaga, J. & Otaegui, D. (2014) Circulating microparticles reflect treatment effects and clinical status in multiple sclerosis. Biomark Med, 8, 653–661.

Sanders, H., Thompson, R.H., Wright, H.P. & Zilkha, K.J. (1968) Further studies on platelet adhesiveness and serum cholesteryl linoleate levels in multiple sclerosis. J Neurol Neurosurg Psychiatry, 31, 321–325.

Schallmoser, K. & Strunk, D. (2013) Generation of a pool of human platelet lysate and efficient use in cell culture. Methods in molecular biology, 946, 349–362.

Semple, J.W., Italiano, J.E., Jr. & Freedman, J. (2011) Platelets and the immune continuum. Nature reviews. Immunology, 11, 264–274.

Sheremata, W.A., Jy, W., Horstman, L.L., Ahn, Y.S., Alexander, J.S. & Minagar, A. (2008) Evidence of platelet activation in multiple sclerosis. Journal of neuroinflammation, 5, 27.

Simon, D.I. (2012) Inflammation and vascular injury: basic discovery to drug development. Circulation journal : official journal of the Japanese Circulation Society, 76, 1811–1818.

Sonia D’Souza, C., Li, Z., Luke Maxwell, D., Trusler, O., Murphy, M., Crewther, S., Peter, K. & Orian, J.M. (2018) Platelets Drive Inflammation and Target Gray Matter and the Retina in Autoimmune-Mediated Encephalomyelitis. Journal of neuropathology and experimental neurology, 77, 567–576.

Steinman, L. (2012) Platelets provide a bounty of potential targets for therapy in multiple sclerosis. Circulation research, 110, 1157–1158.

Tsai, H.H., Niu, J., Munji, R., Davalos, D., Chang, J., Zhang, H., Tien, A.C., Kuo, C.J., Chan, J.R., Daneman, R. & Fancy, S.P. (2016) Oligodendrocyte precursors migrate along vasculature in the developing nervous system. Science, 351, 379–384.

Warnke, P.H., Humpe, A., Strunk, D., Stephens, S., Warnke, F., Wiltfang, J., Schallmoser, K., Alamein, M., Bourke, R., Heiner, P. & Liu, Q. (2013) A clinically-feasible protocol for using human platelet lysate and mesenchymal stem cells in regenerative therapies. Journal of cranio-maxillo-facial surgery : official publication of the European Association for Cranio-Maxillo-Facial Surgery, 41, 153–161.

Woodruff, R.H., Fruttiger, M., Richardson, W.D. & Franklin, R.J.M. (2004) Platelet-derived growth factor regulates oligodendrocyte progenitor numbers in adult CNS and their response following CNS demyelination. Molecular and Cellular Neuroscience, 25, 252–262.

Zhou, Y.X., Flint, N.C., Murtie, J.C., Le, T.Q. & Armstrong, R.C. (2006) Retroviral lineage analysis of fibroblast growth factor receptor signaling in FGF2 inhibition of oligodendrocyte progenitor differentiation. Glia, 54, 578–590.

Zlokovic, B.V. (2008) The blood-brain barrier in health and chronic neurodegenerative disorders. Neuron, 57, 178–201.

