## Supplementary figures and images for "Circulating Platelets Modulate Oligodendrocyte Progenitor Cell Differentiation During Remyelination"

### Supplementary Figure 1

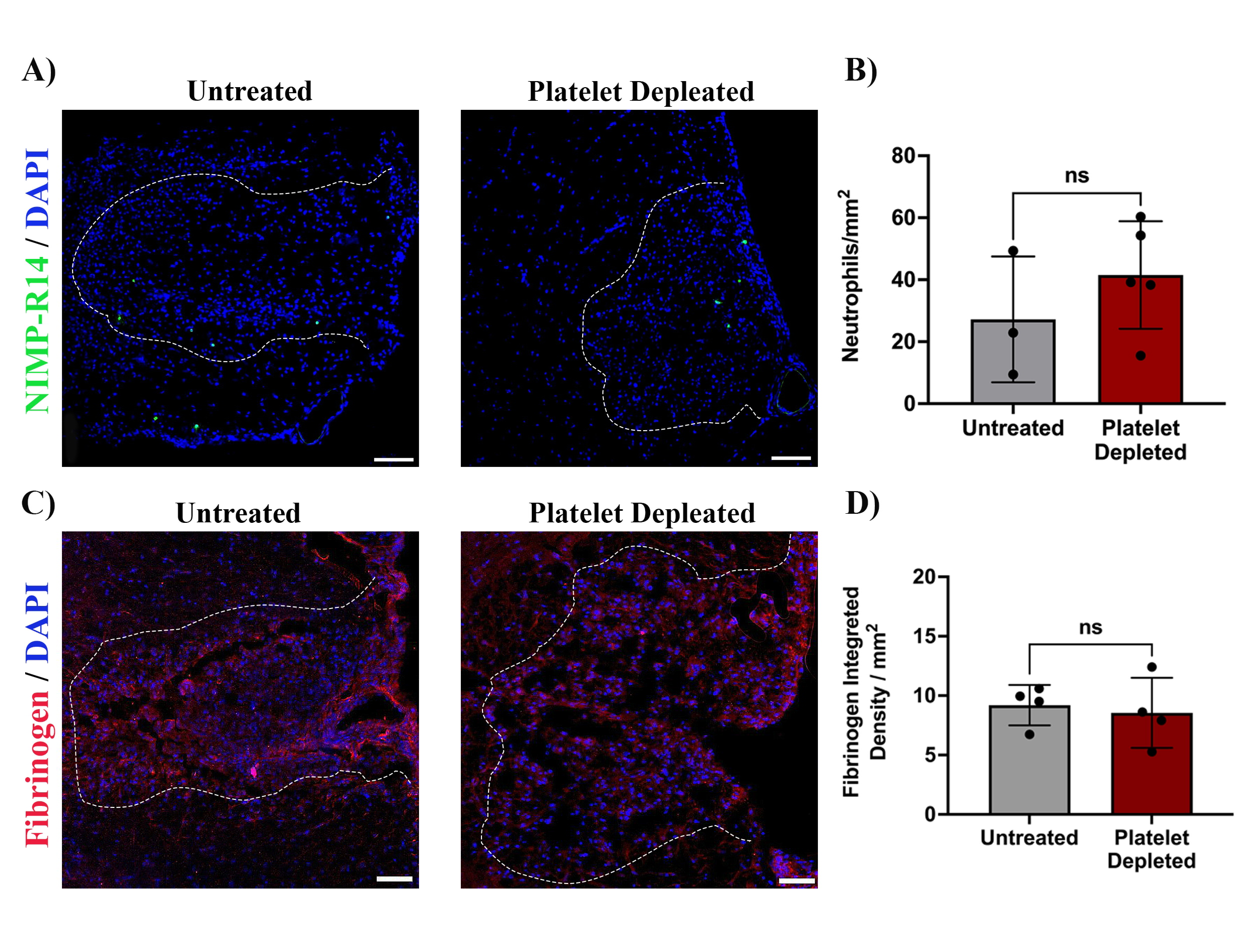

### Supplementary Figure 2

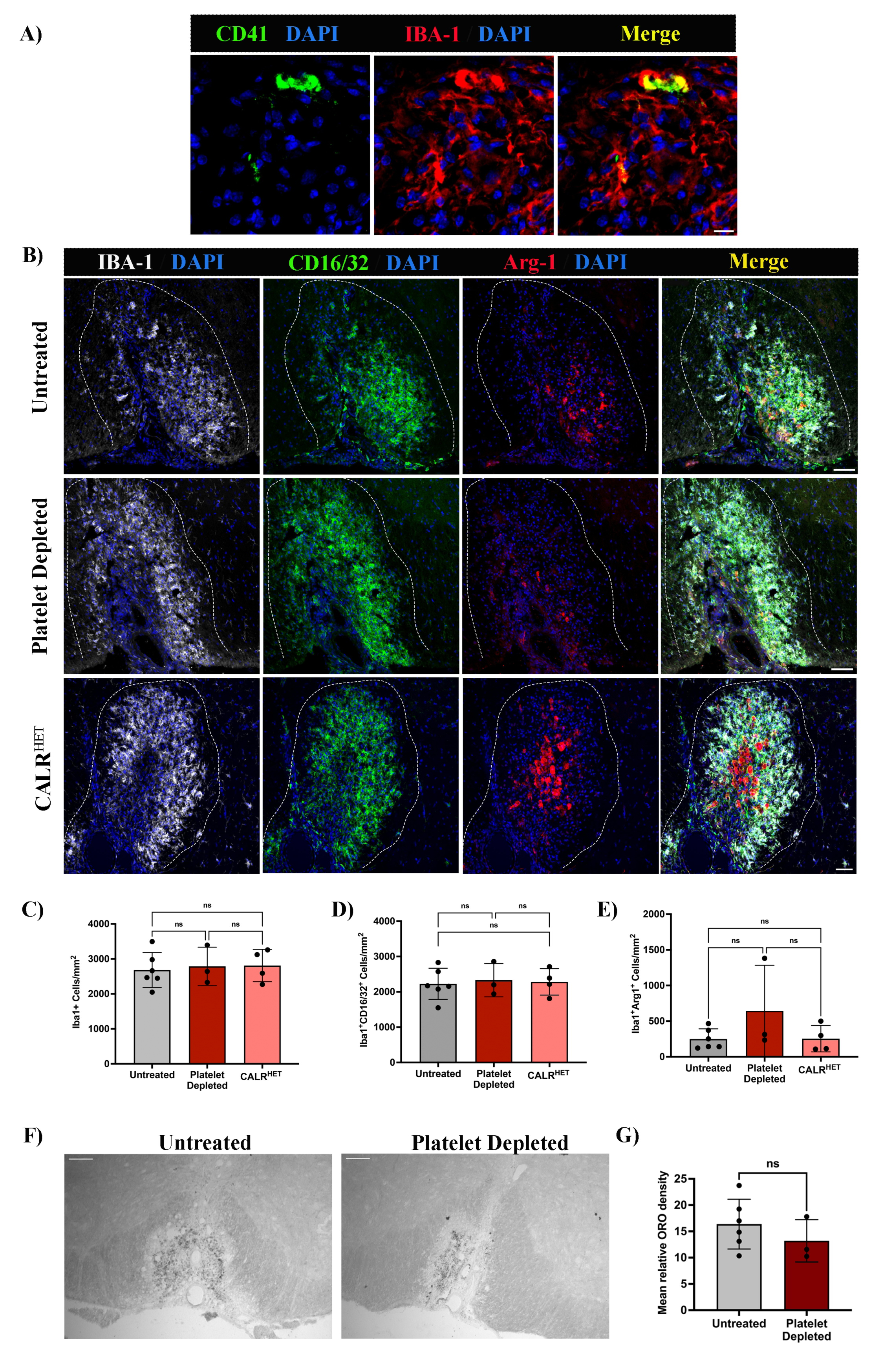
